# Atlantic cod (Gadus morhua) ecotypes, not inversion frequencies, underlie divergence in egg buoyancy distribution

**DOI:** 10.1101/2025.09.28.679046

**Authors:** Rebecca Krohman, Simon Henriksson, Esben Moland Olsen, Halvor Knutsen, Rebekah A. Oomen

**Affiliations:** University of New Brunswick Saint John, 100 Tucker Park Rd, Saint John, NB E2K 5E2; University of Gothenburg, Tjärnö Marine Laboratory, Hättebäcksvägen 7, 452 96 Strömstad, Sweden; Centre for Coastal Research, University of Agder, Campus Kristiansand, Universitetsveien 25, 4630 Kristiansand, Norway; Institute of Marine Research, Flødevigen Research Station, Nye Flødevigveien 20, 4817 His, Norway

**Keywords:** chromosomal inversion, salinity, early life stage, marine fish

## Abstract

Life-history divergence between Atlantic cod (Gadus morhua) ecotypes in Skagerrak is maintained despite gene flow. Fjord and North Sea ecotypes differ in growth, body size, and reproductive ecology. One environmental heterogeneity in Skagerrak is salinity, as coastal habitats experience substantially lower and more variable salinities than offshore waters. Egg buoyancy is highly sensitive to salinity and represents a critical life-history trait shaping survival and recruitment. Inversions on chromosomes 2, 7, and 12 in cod are linked to salinity adaptation and might be important for egg buoyancy and survival. Here, we conducted a buoyancy test on Atlantic cod eggs, spawned from North Sea and Fjord ecotypes, using Coombs apparatus. Genotypes for the inversions on chromosomes 2, 7, and 12 and ecotype were determined in all individuals (N=242). We hypothesized that individuals with different inversion genotypes and ecotypes would have divergent buoyancies. We found that buoyancy was variable and followed a bimodal distribution sorted by ecotype, with Fjord and North Sea individuals having mean buoyancies at neutral salinity (at 7°C) of 27 and 31 ppt, respectively. Therefore, individuals with intermediate buoyancies might be selected against. This finding is consistent with our expectations, as Fjord ecotypes experience lower salinities compared to North Sea ecotypes. Interestingly, there was little evidence that the inversions on chromosomes 2, 7, and 12 impacted buoyancy. Overall, we provide experimental evidence of the relative contributions of ecotypic background and inversions on a key life history trait, which helps to explain the maintained divergence among ecotypes.

## 1 Introduction

Understanding the genetic basis of egg buoyancy in marine fish can be vital for the persistence of fish populations in the future. Many marine fish, including Atlantic cod (*Gadus morhua*), Cape hake (*Merluccius capensis*), and Sardine (*Sardina pilchardus*), spawn neutrally or positively buoyant eggs that grow in the pelagic (Sundby and Kristiansen, 2015). Some northern areas of the Atlantic are experiencing freshening events caused by currents bringing melted sea glacier water south (Cyr and Galbraith, 2021). The density of water decreases as temperature increases, and it increases with salinity. If the density of fish eggs is higher than the density of water, the eggs will sink and die. Therefore, decreasing salinities and increasing temperatures can cause extreme egg die-offs. Understanding the genetic and phenotypic influences on egg buoyancy variation can help make informed predictions on the viability of spawned eggs.

Atlantic cod is an endangered marine fish found across the North Atlantic (Sobel, 1996). Due to differences in life history traits, cod is identified as different ecotypes in the eastern Atlantic and as different populations in the western Atlantic. In the east Atlantic, two ecotypes of cod are known as the Fjord and North Sea ecotypes (Knutsen et al., 2018). These ecotypes spawn in the same areas in coastal Skagerrak and are able to produce hybrids in a laboratory setting (Oomen et al. 2021; Henriksson et al., in prep). However, hybrids are rarely found in the wild (Barth et al., 2019). Hybrid depression, when hybrids have lower survival than non-hybrids, might therefore be present and can contribute to speciation. Environmental variables, including salinity, might be an ecological barrier preventing these ecotypes of cod from spawning interactions (Nissling and Westin, 1997). For example, Eastern Baltic cod spawn in salinities as low as 15 ppt, whereas Skagerrak cod spawn in salinities no lower than 21 ppt (Nissling and Westin, 1997). Therefore, uncovering the buoyancy of the ecotypes and hybridized individuals can provide insights into the lack of occurrence of intermediate genotypes in the wild and into the population structure of cod ecotypes in the Eastern Atlantic.

Cod egg buoyancy is a complex trait closely linked to survival, as healthy eggs maintain neutral buoyancy that allows them to remain suspended in the water column (Craik and Harvey, 1987). Too high buoyancy can cause desiccation at the ocean surface and too low buoyancy can cause sinking into low-oxygen layers. Water density, water movement (e.g., wind), and individual egg buoyancy all interact to determine the depth at which the egg resides (Saborido-Rey, 2003). Furthermore, the specific water column layer at which eggs have neutral buoyancy can assist eggs with being retained in fjord spawning areas or distributed to other areas (Stenevik et al., 2008; Nissling and Vallin, 1996). The hatching depth, directly influenced by buoyancy, is critical for post-hatching survival, shaping access to food and exposure to environmental variables such as predators and oxygen levels (Saborido-Rey, 2003; Nissling and Vallin, 1996). Amino-acid, lipid, inorganic salt, and water content within cod eggs determine buoyancy at spawning, but the osmotic control of water within the egg is what controls buoyancy throughout growth (Craik and Harvey, 1987; Mangor-Jensen, 1987). Therefore, the salinity environment of a cod egg can have major impacts on egg buoyancy and survival to maturity (Knutsen et al., 2007).

Egg buoyancy is dependent on many factors, including population, spawning time, and maternal quality. There are weak correlations between egg size and buoyancy (Nissling and Westin, 1997; Nissling et al., 1994) but strong correlations between spawning time and buoyancy (Saborido-Rey, 2003; Nissling and Vallin, 1996; Jung et al., 2012). Size at hatch, which correlates with egg size, decreases over the spawning season (Roney et al., 2017). One study found that small eggs have moderate buoyancies, and large eggs can have low or high buoyancies depending on spawning time (Kjesbu et al., 1992). Furthermore, buoyancy decreases during development, with an increase just before hatching (Nissling and Vallin, 1996).

Inversions are large-scale mutations in which a genomic segment is flipped in orientation, resulting in suppressed recombination in heterozygotes (Jay et al., 2024; Kirkpatrick, 2010). Inversions can trap one or multiple genes together that contribute to phenotypic traits. For example, inversions contribute to rapid adaptation from marine to freshwater environments in stickleback (*Gasterosteus aculeatus*; Jones et al., 2012). Inversions might also be important for cold water adaptation in Arctic cod (*Arctogadus glacialis*; Hoff et al., 2024). Most evidence for the role of chromosomal inversions in fish phenotypes is correlational, derived from observational or population-genomic studies. Causality is difficult to establish. To move beyond inference, experimental methods are needed to directly test the functional consequence of inversions on ecologically relevant traits (Wellenreuther et al., 2025).

Three inversions in Atlantic cod (inv2, inv7, and inv12) are correlated with salinity in the wild (Sodeland et al., 2016; Berg et al., 2015; Sodeland et al., 2022; Kess et al., 2020) and might influence egg buoyancy. For example, a region of inv12 associated with a double crossover, which is highlighted as a divergent region between North Sea and Fjord ecotypes, includes a vitellogenin gene cluster likely important for egg buoyancy (Matschiner et al., 2022; Henriksson et al., 2023). Inv7 might be important for salinity adaptation (Sinclair-Waters et al., 2018). However, there is no experimental evidence that these inversions impact Atlantic cod egg buoyancy, and thus survival. Understanding the role of the inversions in buoyancy variation can help inform management and conservation that takes into account population structure (Pampoulie et al., 2023; Nielsen et al., 2009).

We conducted an experiment to investigate how inversions and ecotypes contribute to variation in cod egg buoyancy. We hypothesized that mean buoyancy would differ among eggs with different inversion genotypes according to the correlations between inversion frequencies and salinity in the wild. We also hypothesized that Fjord, North Sea, and hybrid ecotype eggs have different mean densities. We predicted that the Fjord ecotype would have a lower density (i.e., floats at lower salinities) than the North Sea ecotype, with hybrids showing intermediate densities.

## 2 Materials and Methods

### 2.1 Spawning population

We used a controlled, semi-natural, outdoor, human-made, flow-through seawater pond located at the Flødevigen Research Station, Institute of Marine Research in Norway. The pond has a diameter of approximately 50 meters, a maximum depth of 4 meters, and flow-through water from 75 meters deep offshore (Figure 1). Water inflow occurs at the bottom of the pond, creating a stable environment in the deeper waters of the pond. The surface of the pond is exposed to the natural environment, creating vertical environmental gradients. Surface water temperatures throughout the year can range from freezing (at -2°C) to 20°C. Surface water salinity decreases when it rains, but at approximately 0.5 meters deep, salinity is in normal ranges of about 34 ppt (RK personal observations).

**Figure 1.**
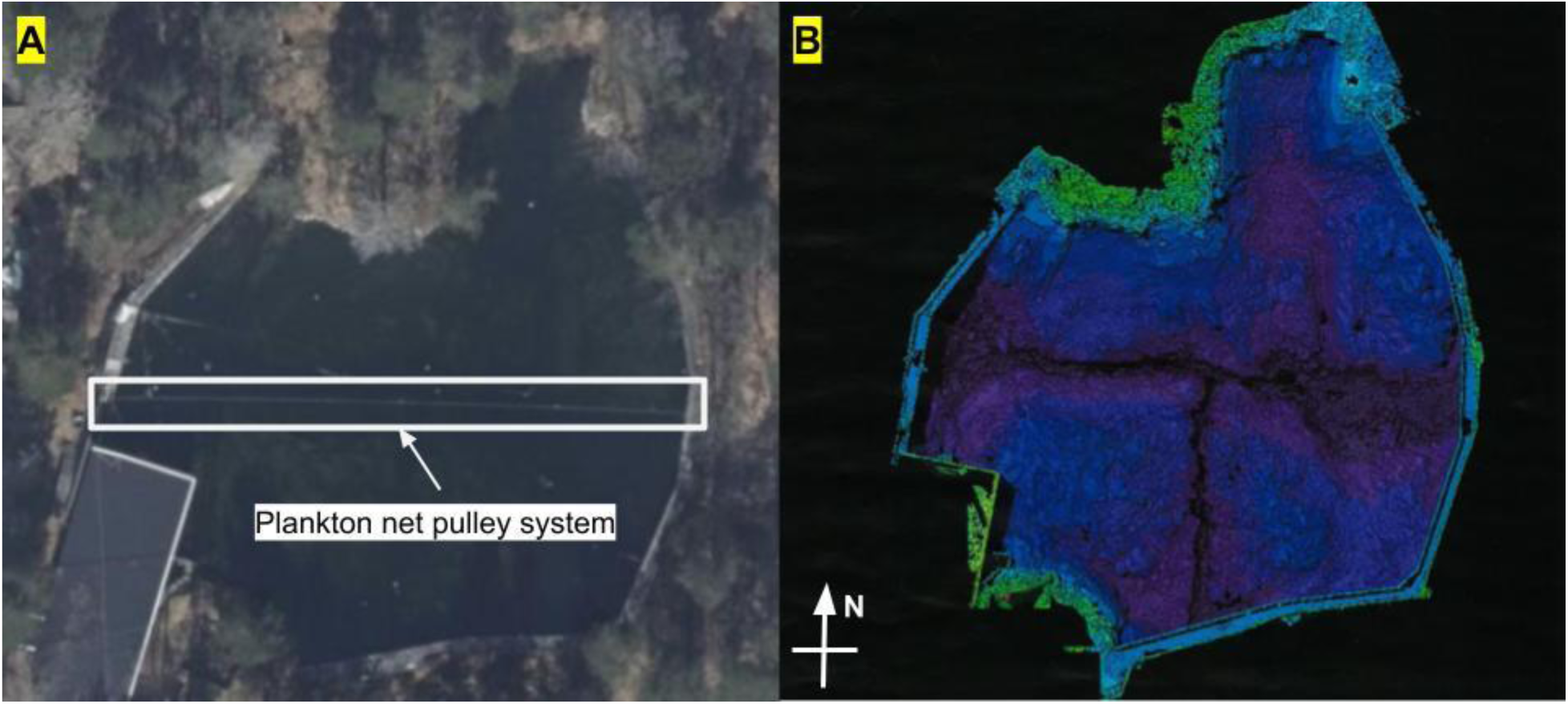
Satellite photo of the cod pond from <u>norgeibilder.no</u> (A) and depth scan of the cod pond (B). The plankton net pulley system set up across the pond is marked in white.

Approximately 50 adult cod inhabit the pond. Individuals from the North Sea and Fjord ecotypes were collected from the southern coast of Norway (N=45: 17 from approximately 58.4° latitude 8.8° longitude; 28 from approximately 58.3° latitude and 8.6° longitude) in Fall 2022 and allowed to spawn freely in the pond in winter 2022 before being released back into the wild. The F1 population freely spawned the F2 generation in Winter 2024 and 2025. The F0 generation was polymorphic for inv2, inv7, and inv12, and it is likely that both the F1 and F2 generations are polymorphic as well. This population is ideal for testing phenotypic correlations with inv2, inv7, and inv12 frequencies.

### 2.2 Egg collection and holding

Before eggs were collected, the temperature, salinity, and dissolved oxygen of the pond were measured at one meter depth using a YSI ProSolo meter with an ODO/CT probe. The average (±standard error (SE)) pond temperature, salinity, and dissolved oxygen were 4.95 (±1.044), 33.58 (±0.20), and 107.03 (±8.68), respectively. We used a plankton pulley net (320 μm mesh size and 46 cm diameter) system that sits at the surface of the pond across the deepest part of the pond (Figure 1) to collect eggs. Eggs were collected in four batches numbered 2, 3, 4, and 5, on February 17th and 25th and March 3rd and 10th, 2025, respectively. To collect eggs, we pulled the plankton net across the pond twice by pulling the net about 10 m/min in one direction and about 16.7 m/min in the opposite direction. Different pulling speeds allowed us to collect a variety of individuals with different buoyancies. At either end of the pond, eggs were gently removed from the collection container into a bucket containing surface pond water. The water in the bucket was tested for pH, ammonia, nitrate, and nitrite (API saltwater master test kit), with average (±SE) measurements being 7.95 (±0.03), 0 (±0), 0 (±0), and 0 (±0), respectively. There were changes made to the above collection protocol in batches 3 and 4 (resulting in batch 4b) that can be found in Appendix A.

The eggs were sorted using a plastic pipette (4 mm inner diameter) into early (body length <50% of the egg) and late stages (body length >50% of the egg) using a Leica ES2 dissecting microscope. Individuals were placed, separated by stage, into one of two circular (19 cm inside edge diameter) holding tanks with a 200 μm mesh bottom and low surface water turnover (Figure 2). The water in the holding tank was from 75 m deep offshore and macrofiltered with a 40 μm filter. The average (±SE) temperature, salinity, and pH of the holding tanks were 7.1 (±0.29)°C, 33.7 (±1.10) ppt, and 8.0 (±0.03), respectively. Individuals in the holding tank were exposed to 210 lux light, measured with a light meter Art 15-341 (Biltema), for 10 hours per day.

**Figure 2.**
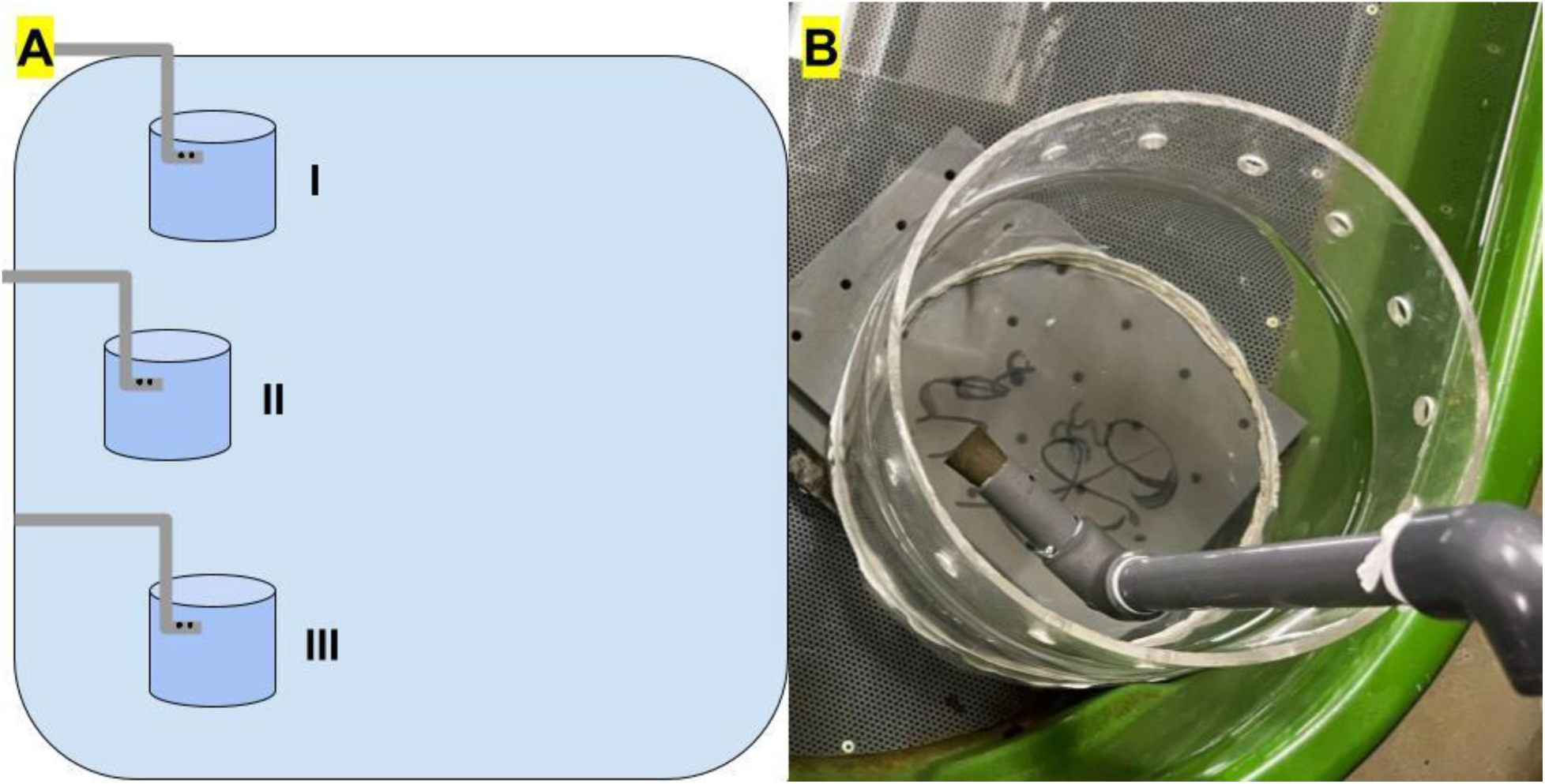
A schematic of the setup of the holding tank and holding containers I, II, and III, in which individuals were held before experimentation (A) and a close-up of one of the holding tanks (B). The schematic shows the tubing that the water flowed through (grey), the bore holes at the end of the tubing that helped control water flow (black), and the small circular holding containers (dark blue) inside the large holding tank (light blue). Note the mesh at the bottom of the holding container in part B. The grey blocks below the holding container were used to prop the holding container higher so that the holes on the top of the container would be above the water line to prevent any eggs from escaping through the holes.

### 2.3 Buoyancy experiment

We determined the buoyancy of cod eggs using a density-gradient column (Coombs apparatus). Low salinity water (average (±SE) = 15.3 (±0.02) ppt) was created by mixing water from a spout in the holding tank with freshwater filtered by a Synergy UV-R system filter, at a calculated ratio to get 15 ppt. Water salinity was measured using a YSI ProSolo meter with an ODO/CT probe (±1.0% of reading) before being placed in the fill container. Water added to the tank was at ambient water and lab temperature (∼ 7°C). All trials took place at least 1.25 hours and no more than 120 hours after collection. Once the salinity column was filled, 4 beads with known densities (yellow, olive, red, and clear bead densities were 1.0265, 1.0227, 1.0170, and 1.0128 g/cm^3^, respectively; Martin Instrument Co.) were placed in the salinity column (see Supplementary Figure 1). Then, we placed a maximum of four eggs in the column using a pipette. We used individuals from each stage because there are identifiable differences between stages, allowing us to identify them after they were removed from the column. For example, stages 1, 2, 3, and 4 were identified by the following characteristics: a visible embryo, no visible embryo, a visible body, and visible eyes, respectively. After the eggs stopped sinking, we measured the top of each bead. Then we found, staged, and measured the location of the eggs. To remove the beads and eggs from the column, the water in the column was carefully poured through a 5cm inner-diameter circular container with a 200 μm mesh bottom. All eggs were sampled for DNA extraction purposes. A total of 115 tank replicates were run, and 250 individual eggs were measured (n = 48, 63, 28, 35, and 76 for batches 2, 3, 4, 4b, and 5, respectively).

Egg density (g/cm^3^) was estimated based on the locations of the beads in the salinity column. The red and olive bead densities and heights were used to calibrate the relationship between depth and density. The density of water at 7°C with different salinities was taken from omnicalculator.com/physics/water-density to relate buoyancy to neutral salinities. Eggs were classified as having neutral buoyancy at a given salinity if the egg density was less than or equal to the water density at 7°C.

### 2.5 Genetic analyses

All eggs were fixed in 96% ethanol and held in the lab for a maximum of 36 hours before being placed into a freezer at -20°C for holding. Eggs (n=242) were sent via overnight mail to IdentiGEN LTD, Dublin, Ireland, for genotyping of 4000 single-nucleotide polymorphisms (SNPs; Shapero, 2013). The genetic panel was originally designed for genetic monitoring of Scandinavian cod populations (Henriksson et al., in prep; *cf.* Andersson et al., 2024). SNP data for 3561 loci were received from 230 individuals. In brief, ecotypes, inversion genotypes, and parental relationships were assigned to eggs using assignPOP (v1.3.0; Chen, 2024), PCAs, and COLONY (v2.0.7.1; Wang, 2004), respectively. Further information is found in the supplementary methods.

To explore genomic differences between the ecotypes, eggs were grouped according to their majority assignment in assignPOP (see Supplementary methods). Locus-wise *F*_ST_ between the two ecotypes was calculated using the strataG package (v2.5.1; Archer et al., 2016). Based on the modality of the buoyancy results (see section 3.3), we performed a similar calculation of *F*_ST_ between eggs of “low” and “high” buoyancy. The cutoff between the two buoyancy groups was defined as the midpoint between the two modes (Figure 3), which were defined using a custom script. Outlier analysis for both pairwise comparisons was performed with the OutFLANK package (v0.2; Whitlock and Lotterhos, 2014), using default settings. Loci displaying significant divergence between groups were explored for gene ontology (GO) terms on the PANTHER database (Thomas et al., 2022), and suggested gene functions in fish were explored by searching for papers on Google Scholar (search string: “*gene name* fish”).

**Figure 3.**
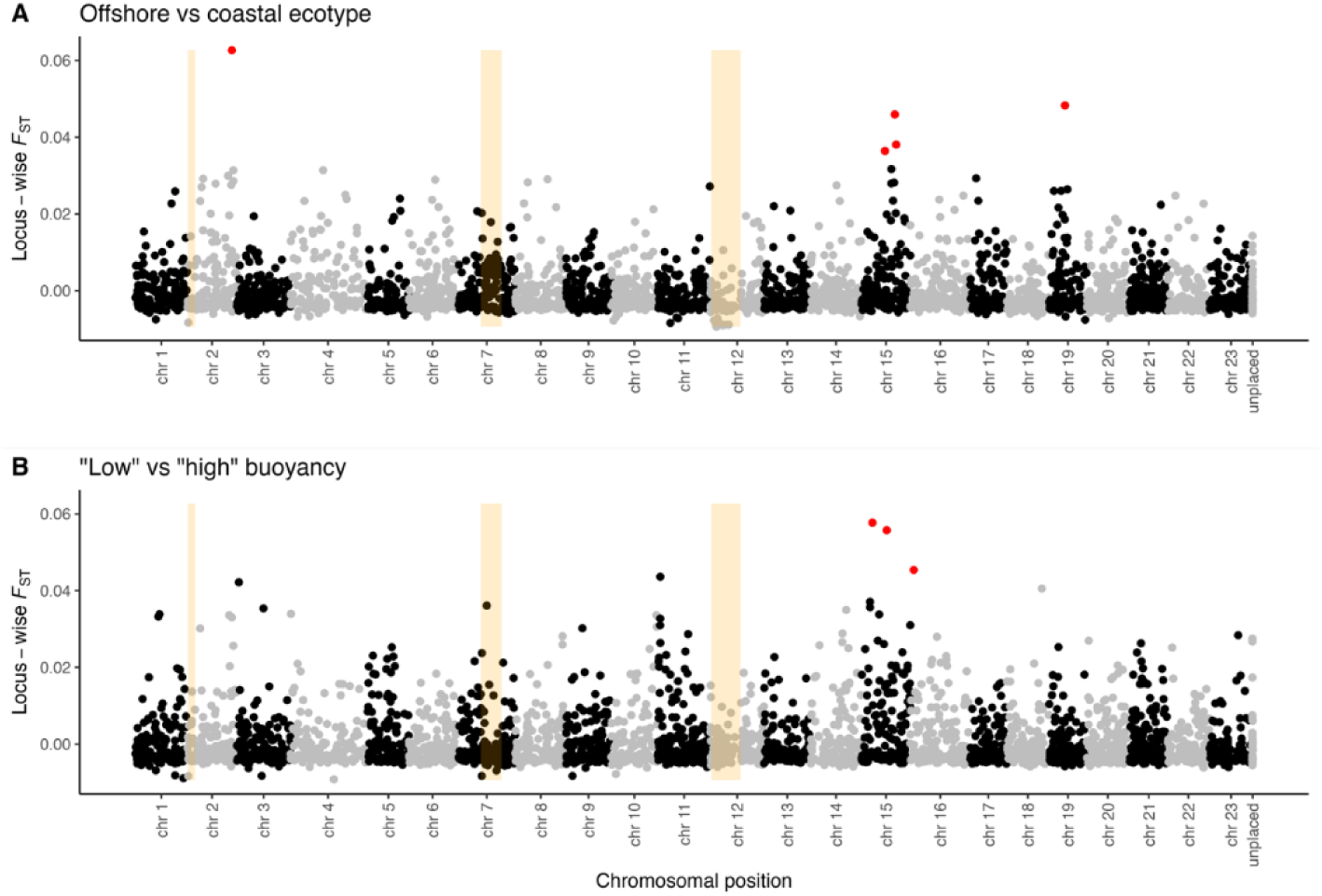
Pairwise F_ST_ between North Sea and Fjord ecotype eggs (A), and “low” and “high” buoyancy (B). Inverted regions on chromosomes 2, 7, and 12 are highlighted in orange, and outlier loci from each pairwise comparison are indicated with red points.

### 2.6 Statistical methods

A generalized linear mixed model using template model builder (glmmTMB) was created to test the hypothesis that inversions and ecotypes impact egg buoyancy via the glmmTMB function in the glmmTMB package (v1.1.11; Brooks et al., 2017). The three inversions, inv2, inv7, and inv12, were factors (3, 2, and 2 levels for inv2, inv7, and inv12, respectively) tested as main effects along with the probability of belonging to the North Sea ecotype (Equation 1). Egg stage, parent 1 and 2 ID, time since collection, and batch were included as random effects in the model originally, but were dropped due to minimal variation being explained by each effect (see Supplementary methods). The inv2 heterozygote genotype was used as a reference level. To test model fit, the residuals were visually checked in an x-y plot, histogram, and qq-plot. There were no major deviations of the residuals. The model output was predicted with the ggpredict function in the ggeffects package (v2.2.1; Lüdecke, 2018) for graphing purposes.

All statistics were calculated in R Studio v4.4.2 (R Core Team, 2023), and figures were created in GraphPad Prism (v101) and ggplot2 (Wickham, 2016). An alpha value of 0.05 was used throughout.

Equation 1. Egg buoyancy ∼ inv2 + inv7 + inv12 + probability of belonging to the North Sea ecotype

## 3 Results

### 3.1 Variability of egg buoyancy

All eggs had a neutral buoyancy between salinities of 22 and 35 ppt. The number of individuals with neutral buoyancy at a given salinity followed a bimodal distribution with peaks forming around 26-27 ppt and 32-33 ppt (Figure 4). The lowest and highest measured buoyancies (-Density (g/cm^3^)) were -1.0274 and -1.0178, respectively. The standard deviation and error of the buoyancy of all eggs were 0.00238 and 0.00015, respectively.

**Figure 4.**
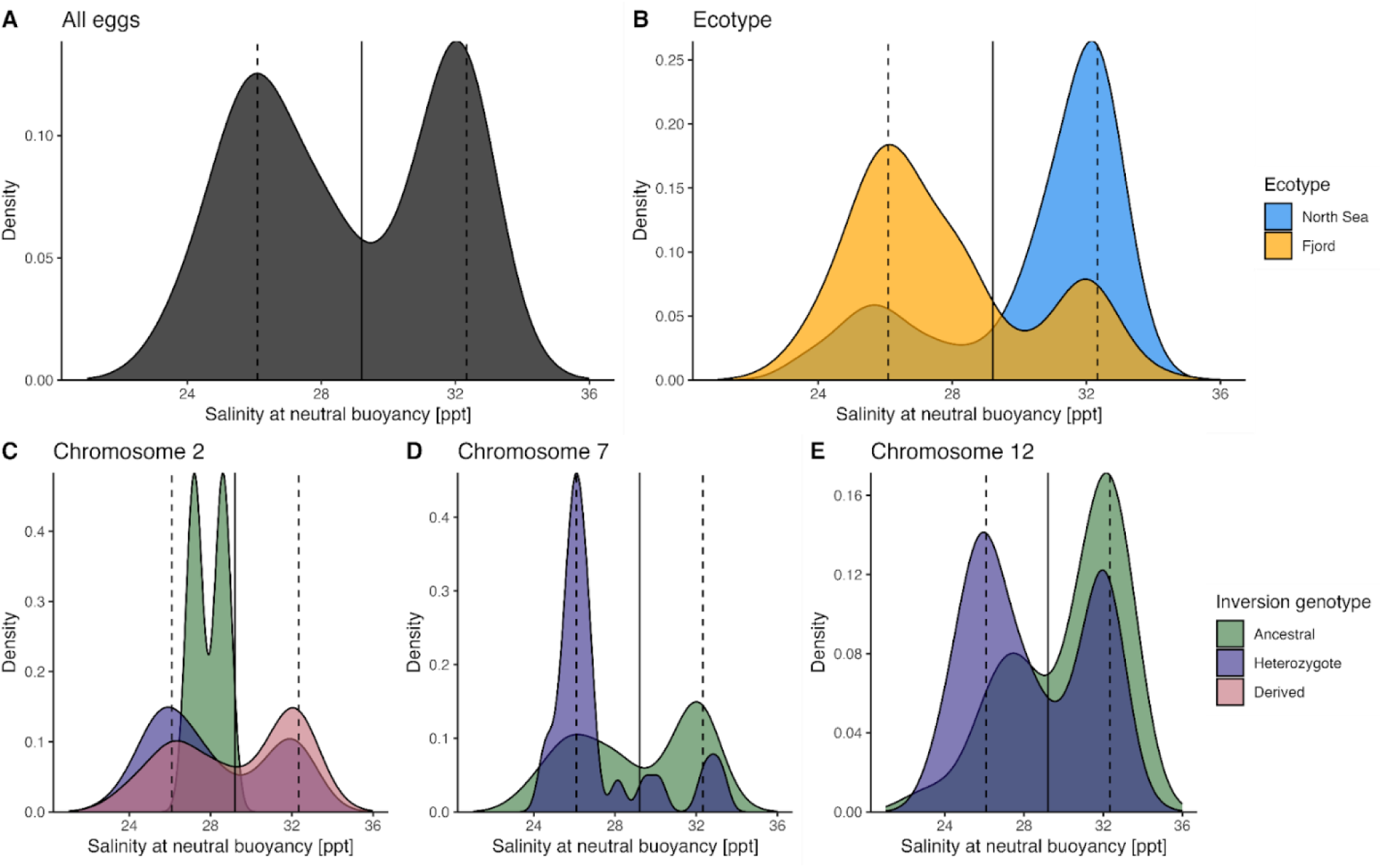
Salinity at neutral buoyancy at 7°C for all eggs (A), and grouped by ecotype (B), or genotype at inv2 (B), inv7 (C), and inv12 (D). The dashed lines show the two buoyancy modes inferred, and the solid line shows the cutoff value used to distinguish eggs with “low” and “high” buoyancy. Note the differences in the y-axis range between panels.

### 3.2 Effect of genomic variables on buoyancy

In histograms sorted by inversion genotypes, there is no clear division between inversion genotypes for inv2, inv7, and inv12, but there is a division based on ecotype (Figure 4). The mean buoyancy of the inv7 heterozygote was marginally significantly different from the inv7 derived genotype, while inv2 and inv12 genotypes did not significantly impact buoyancy (Table 1; Figure 5). However, ecotype significantly impacted buoyancy (Table 1; Figure 5). The predicted line of best fit found a slightly negative relationship between buoyancy and belonging to the North Sea ecotype (Figure 5). The model fit with an R^2^ of 0.199.

**Figure 5.**
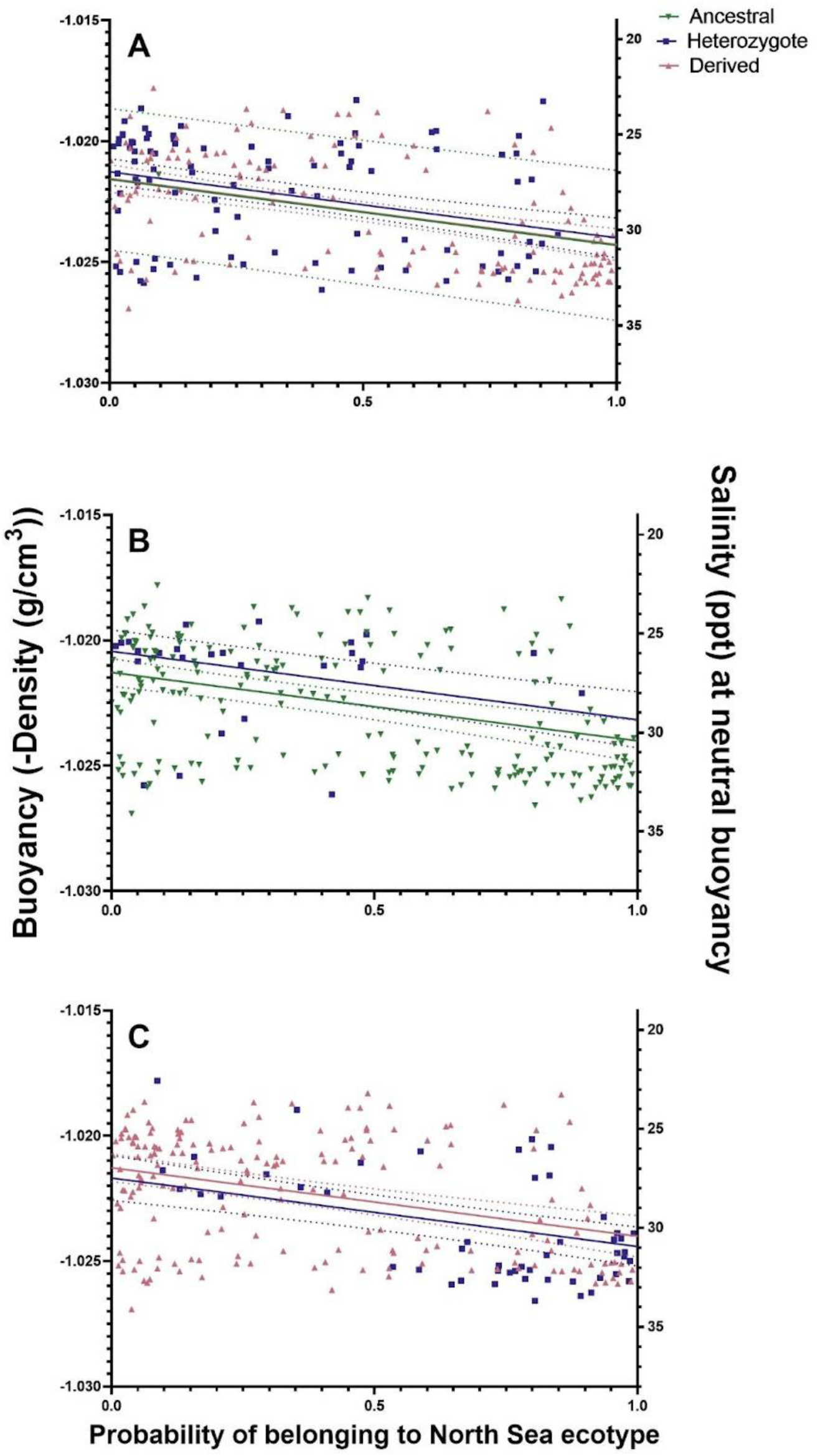
The buoyancy (-Density (g/cm^3^)) of eggs is dependent on the probability of belonging to the North Sea ecotype but not inversion 2 (A), 7 (B), or 12 (C) genotypes. The ancestral, heterozygote, and derived genotypes are represented by green inverted triangles, blue squares, and pink triangles, respectively. The neutral buoyancy at a given salinity (ppt) at 7°C is provided to help link salinity with buoyancy. Plotted points are the raw data points. The dashed lines display the 2.5% to 97.5% confidence interval for the line of best fit (from the created model). N=230.

**Table 1.** The model result for the effect of inversion genotypes on buoyancy (Equation 1) includes the predictors, estimates, confidence intervals, and p-values.

| Effect of inversions and ecotype on egg buoyancy |  |  |  |  |
| --- | --- | --- | --- | --- |
| *Predictors* | *Estimates* | *Standard error* | *Z value* | *p-value* |
| (Intercept) | -1.0213 | 0.0002 | -3632 | <0.001 |
| Inv2 Ancestral | -0.0003 | 0.0015 | 0 | 0.838 |
| Inv2 Derived | -0.0003 | 0.0003 | -1 | 0.348 |
| Inv7 Heterozygote | 0.0008 | 0.0004 | 2 | 0.061 |
| Inv12 Heterozygote | -0.0004 | 0.0004 | -1 | 0.275 |
| Probability of belonging to the North Sea ecotype | -0.0027 | 0.0005 | -6 | **<0.001** |
| Observations | 230 |  |  |  |
| $R^2$ / $R^2$ adjusted | 0.220 / 0.199 | | | |

Bolded values indicate significance.

### 3.3 Genomic differentiation

Genetic differentiation between North Sea and Fjord cod was heterogeneous across the genome (Figure 3). In contrast to previous studies, however, this divergence was not strongly associated with chromosomal inversions. Five outlier loci were detected on chromosomes 3 (n=1), 15 (n=3), and 19 (n=1). All were located within annotated genes in the GadMor3 genome, with suggested functional roles in neurogeneration, growth, gastrulation, environmental adaptation, sexual maturation, and muscle filament assembly in fish (see Supplementary Notes). Similarly, genetic differentiation between eggs of “low” and “high” buoyancy was associated with regions outside of the inversions (Figure 3). Three outlier loci on chromosomes 15 (n=2), and 16 (n=1) were detected, two within annotated genes, and one located between two genes. The genes are involved in growth, fin development, electric synapse function, response to viral infection and DNA damage (see Supplementary Notes). Neither ecotypes nor buoyancy groups showed signs of divergence at loci within the region of double crossover inside the chromosome 12 inversion (see Supplementary Figure 2).

### 3.4 Random effects on buoyancy

Egg buoyancy increased with increasing time since collection, with mean (±SE) buoyancy increasing from -1.0230 (±0.00074) before 1 hour to -1.0235 (±0.00048) after 100 hours (see Supplementary Figure 3). Egg buoyancy between early (-1.0230 ±0.00025) and late (-1.0225 ±0.00019) stage individuals was similar (see Supplementary Figure 4). Egg buoyancy in batches 2 and 4b is higher than all other batches, and batch 4 is lower than all batches except batch 3 (see Supplementary Figure 5). Mean (±SE) egg buoyancies were -1.0217 (±0.00035), - 1.0237 (±0.00029), -1.0245 (±0.00019), -1.0213 (±0.00022), and -1.0227 (±0.00028) in batches 2, 3, 4, 4b, and 5, respectively. There were differences in egg buoyancy between parents 1 and 2 individuals (see Supplementary Figure 6).

## 4 Discussion

In our study, cod eggs showed large variability in buoyancy, ranging from -1.0274 to - 1.0178, respectively. Ecotype had a significant effect on buoyancy, whereas inversions on chromosomes 2 (inv2), 7 (inv7), and 12 (inv12) did not. Because ecotype captures broader genomic variation, it likely integrates multiple genetic factors shaping buoyancy. When considering other factors, buoyancy varied with time-since-collection and among batches and parents, though not among egg developmental stages.

### 4.1 Variability of buoyancy

Cod eggs had a salinity at neutral buoyancy spanning from 23 to 35 ppt, indicating that this is a variable trait. We found a larger salinity range in this experiment, on southern Norwegian cod, compared to northern Norwegian cod, which are found to have a buoyancy ranging between 30 and 34 ppt (Stenevik et al., 2008). Salinity at neutral buoyancy had a bimodal distribution, with peaks forming at 26 and 32 ppt. This suggests that a variable with two levels influences buoyancy variation. Furthermore, the bimodal relationship suggests that there is either current or historical selection against individuals whose neutral buoyancy is between 28 and 31 ppt.

### 4.2 Genomic impacts on egg buoyancy

We found that buoyancy decreased with an increasing degree of belonging to the North Sea ecotype, suggesting local adaptation to salinity between ecotypes. North Sea individuals are expected to have a lower buoyancy than Fjord individuals because salinity increases away from the coast, and eggs would need to have a lower neutral buoyancy. Previous studies on cod in Norway and Sweden also found that ecotype impacts egg buoyancy, generally with neutral buoyancies at salinities that the population is normally exposed to (Nissling and Westin, 1997; Nissling and Vallin, 1996). Differences in egg buoyancy are suggested to be a physical barrier between cod ecotypes that spawn in the same areas, creating a post-spawning, pre-zygotic barrier (Stenevik et al., 2008). Adaptation to local salinities can put cod at higher risk of detrimental climate change impacts and could make intraspecific diversity in salinity tolerance particularly important for population survival.

The observation that inv2, inv7, and inv12 did not impact buoyancy was surprising because: (1) inv2 is strongly correlated with salinity in the wild (Berg et al., 2015), (2) genes within inv7 are linked to low salinity adaptation (Sodeland et al., 2022; Sinclair-Waters et al., 2018), and (3) genes found in inv12 are proposed to control egg buoyancy (Matschiner et al., 2022). Although these inversions do not significantly impact egg buoyancy, they might play a role in salinity adaptation at other life stages or help egg survival in low-salinity conditions.

Our results on the role of inversions in buoyancy display the importance of experimental tests in genomic research to compare with trends found with observational data. We show a disconnect between observational and experimental studies of chromosomal inversions in Atlantic cod. For instance, the vitellogenin gene cluster located within the inversion on chromosome 12 was previously suggested to impact buoyancy in eggs (Matschiner et al., 2022), but here we see no evidence of this. Experiments complement observational studies to fully understand the role of genomic features in environmental adaptation.

### 4.3 The impact of other factors on buoyancy

There was little difference in buoyancy between early and late stage individuals, contrary to previous research (Nissling and Vallin, 1996; Jung et al., 2012; Saborido-Rey, 2003).

Buoyancy increased with the amount of time individuals spent in the lab, which correlates with development. Interestingly, multiple studies found that cod buoyancy generally changes to be more similar to the salinity (Schmidt et al., 2024; Nissling and Vallin, 1996), possibly because osmosis into the oocyte membrane would equalize egg buoyancy (Fridman, 2020). However, we found the opposite trend, indicating that the relationship between egg buoyancy and time since collection was likely caused by a change in lifestage. As individuals spent more time in the lab, they grew older. Previous research found that buoyancy increases right before hatching (Jung et al., 2012; Saborido-Rey, 2003).

Buoyancy varied with batch. Batches 3 and 4 might have had lower buoyancies due to collection conditions. Specifically, there was heavy rain during the collection of batch 3, and the top 0.5 m of the pond was fresher than ambient conditions; those with higher buoyancies might have died at the top of the pond. Similarly, in batch 4, individuals with higher buoyancies were blown to the corner of the pond (batch 4b). Differences between batches 2 and 5 might be due to the fact that egg buoyancy generally decreases throughout the spawning season, even within the same female (Kjesbu et al., 1992; Nissling et al., 1994; Saborido-Rey, 2003). Changing egg buoyancy throughout the spawning season can allow repeat spawners to spawn eggs with different vertical distributions, impacting egg distribution and allowing for eggs to hatch at different survival conditions (Kjesbu et al., 1992; Nissling and Vallin, 1996).

We observed differences in buoyancy between the offspring of different parents, consistent with previous research, which found that buoyancy varied across family groups (Jung et al., 2012). Previous studies found that female physical features are not significantly correlated with egg buoyancy but impact larval buoyancy (Jung et al., 2012; Saborido-Rey, 2003; Schmidt et al., 2024). Therefore, observed differences in egg buoyancy between parents might be due to genomic effects, like ecotype, rather than parental effects in cod eggs. Epigenetic effects, parental effects, and genetic variation are challenging to disentangle (Curley et al., 2011).

High variability of buoyancy might be advantageous, as it promotes some egg survival in unpredictable environments. In changing climates, such variability represents a key life-history trait, allowing some individuals to survive even when conditions deviate from the optimum. This is supported by previous research that found the ability of eggs to maintain neutral buoyancy varies between years (Nissling et al., 1994). Another study highlighted the importance of high buoyancy variability across different families and individuals (Jung et al., 2012). Therefore, ecotype is likely one of several factors contributing to egg buoyancy variation.

### 4.4 Conclusion

Cod egg buoyancy is a variable trait strongly influenced by ecotype. The bimodal distribution of egg buoyancy shows that ecotypes differ in buoyancy and that selection might act against eggs with intermediate buoyancies. This suggests adaptation of ecotypes to local salinity conditions. Furthermore, our results that inversions do not contribute substantially to buoyancy variation show the importance of validating correlations observed in the field with experiments. Specifically, common garden experiments combined with statistical modelling can help separate the effects of inversions and of other parts of the genome on physiological traits, like buoyancy (Mérot et al., 2020). Overall, variability in egg buoyancy and its link to ecotype divergence can inform our understanding of the persistence of cod populations under future environmental change.

## Acknowledgements

We respectfully acknowledge the research conducted at the University of New Brunswick, SJ was conducted on unsurrendered and unceded traditional Wəlastəkwiyik (Wolastoqiyik) and Mi’kmaq (part of the Wabanaki) land. This territory is part of the Peace and Friendship Treaties, which did not involve the surrender of lands, waters, or resources and established an ongoing relationship of peace, friendship and mutual respect between equal nations. Tusen takk til Stian Stiansen och Jan Henrickson Simonsen for hjelpen med eksperimentene. Individuals at the Tjärnö Marine Laboratory and Flødevigen Research Station, Institute of Marine Research, particularly Hanne Sannæs and Heidi Fiskaa, are thanked for all their help. Thanks go out to Dr. Scott Pavey and Dr. Alex Zimmer at UNBSJ for their guidance on the project. We’d like to thank Claudia Lacroix, Abigail K. Scher, and Stephanie Maheux for the feedback that they provided.

We have approval from the Animal Ethical Committee of the Norwegian Food Safety Authority (Mattilsynet) to carry out the work with the utmost care for environmental impacts and fish welfare (FOTS ID 28408).

## Data Availability

We have deposited the primary data underlying these analyses in the Mendeley data depository doi: 10.17632/j8kdg9hsc5.1

## Contributions

RK: Conceptualization; Data curation; Formal analysis; Funding acquisition; Investigation; Methodology; Project administration; Visualization; Writing - original draft; Writing - review & editing

SH: Data curation; Formal analysis; Investigation; Methodology; Visualization; Writing - original draft; Writing - review & editing

EMO: Resources; Writing - review & editing

HK: Writing - review & editing

RAO: Conceptualization; Funding acquisition; Methodology; Resources; Supervision; Writing - review & editing

## References

Andersson, L., Bekkevold, D., Berg, F., Farrell, E. D., Felkel, S., Ferreira, M. S., Fuentes-Pardo, A. P., Goodall, J. and Pettersson, M. (2024). How fish population genomics can promote sustainable fisheries: A road map. Annu. Rev. Anim. Biosci. 12, 1–20. 10.1146/annurev-animal-021122-102933

Archer, F. I., Adams, P. E., and Schneiders, B. B. (2017). stratag: An r package for manipulating, summarizing and analysing population genetic data. Mol. Ecol. Resour. 17(1), 5–11. 10.1111/1755-0998.12559

Badyaev, A. V. (2008). Maternal effects as generators of evolutionary change: a reassessment. Ann. N. Y. Acad. Sci. 1133, 151–161. 10.1196/annals.1438.009

Barth, J. M. I., Villegas-Ríos, D., Freitas, C., Moland, E., Star, B., André, C., Knutsen, H., Bradbury, I., Dierking, J., Petereit, C., et al. (2019). Disentangling structural genomic and behavioural barriers in a sea of connectivity. Mol. Ecol. 28, 1394–1411. 10.1111/mec.15010

Berg, P. R., Jentoft, S., Star, B., Ring, K. H., Knutsen, H., Lien, S., Jakobsen, K. S. and André, C. (2015). Adaptation to low salinity promotes genomic divergence in Atlantic cod (*Gadus morhua* L.). Genome Biol. Evol. 7, 1644–1663. 10.1093/gbe/evv093

Brooks, M., Kristensen, K., Benthem, K. van, Magnusson, A., Berg, C., Nielsen, A., Skaug, H., Mächler, M. and Bolker, B. (2017). GlmmTMB balances speed and flexibility among packages for zero-inflated generalized linear mixed modeling. R J. 9, 378.

Catanach, A., Crowhurst, R., Deng, C., David, C., Bernatchez, L. and Wellenreuther, M. (2019). The genomic pool of standing structural variation outnumbers single nucleotide polymorphism by threefold in the marine teleost *Chrysophrys auratus*. Mol. Ecol. 28, 1210– 1223. 10.1111/mec.15051

Chen K, Marschall EA, Sovic MG, Fries AC, Gibbs HL, Ludsin SA (2024). assignPOP: Population Assignment using Genetic, Non-Genetic or Integrated Data in a Machine Learning Framework. R package version 1.3.0.

Craik, J. C. A. and Harvey, S. M. (1987). The causes of buoyancy in eggs of marine teleosts. J. Mar. Biol. Assoc. U. K. 67, 169–182. 10.1017/S0025315400026436

Curley, J. P., Mashoodh, R. and Champagne, F. A. (2011). Epigenetics and the origins of paternal effects. Horm. Behav. 59, 306–314. 10.1016/j.yhbeh.2010.06.018

Cyr, F. and Galbraith, P. S. (2021). A climate index for the Newfoundland and Labrador shelf. *Earth Syst*. Sci. Data 13, 1807–1828. 10.5194/essd-13-1807-2021

Dray, S. and Dufour, A.-B. (2007). Theade4Package: Implementing the duality diagram for ecologists. J. Stat. Softw. 22. 10.18637/jss.v022.i04

Fridman, S. (2020). Ontogeny of the osmoregulatory capacity of teleosts and the role of ionocytes. Front. Mar. Sci. 7. 10.3389/fmars.2020.00709

Gruber, B., Unmack, P. J., Berry, O. F. and Georges, A. (2018). dartr: Anrpackage to facilitate analysis of SNP data generated from reduced representation genome sequencing. Mol. Ecol. Resour. 18, 691–699. 10.1111/1755-0998.12745

Henriksson, S., Pereyra, R. T., Sodeland, M., Ortega-Martinez, O., Knutsen, H., Wennhage, H., and André, C. (2023). Mixed origin of juvenile Atlantic cod (*Gadus morhua*) along the Swedish west coast. ICES J. Mar. Sci. 80(1), 145–157. 10.1093/icesjms/fsac220

Hoff, S. N. K., Maurstad, M., Tørresen, O. K., Berg, P. R., Præbel, K., Jakobsen, K. S. and Jentoft, S. (2024). Chromosomal fusions and large-scale inversions are key features for adaptation in Arctic codfish species. bioRxiv. 10.1101/2024.06.28.599280

IUCN (1996). Gadus morhua: Sobel, J. IUCN Red List of Threatened Species.

Jay, P., Aubier, T. G. and Joron, M. (2024). The interplay of local adaptation and gene flow may lead to the formation of supergenes. Mol. Ecol. 33, e17297. 10.1111/mec.17297

Jones, F. C., Grabherr, M. G., Chan, Y. F., Russell, P., Mauceli, E., Johnson, J., Swofford, R., Pirun, M., Zody, M. C., White, S., et al. (2012). The genomic basis of adaptive evolution in threespine sticklebacks. Nature 484, 55–61. 10.1038/nature10944

Jung, K.-M., Folkvord, A., Kjesbu, O. S., Agnalt, A. L., Thorsen, A. and Sundby, S. (2012). Egg buoyancy variability in local populations of Atlantic cod (*Gadus morhua*). Mar. Biol. 159, 1969–1980. 10.1007/s00227-012-1984-8

Kess, T., Bentzen, P., Lehnert, S. J., Sylvester, E. V. A., Lien, S., Kent, M. P., Sinclair-Waters, M., Morris, C., Wringe, B., Fairweather, R., et al. (2020). Modular chromosome rearrangements reveal parallel and nonparallel adaptation in a marine fish. Ecol. Evol. 10, 638– 653. 10.1002/ece3.5828

Kirkpatrick, M. (2010). How and why chromosome inversions evolve. PLoS Biol. 8, e1000501. 10.1371/journal.pbio.1000501

Kjesbu, O. S., Kryvi, H., Sundby, S. and Solemdal, P. (1992). Buoyancy variations in eggs of Atlantic cod (*Gadus morhua* L.) in relation to chorion thickness and egg size: theory and observations. J. Fish Biol. 41, 581–599. 10.1111/j.1095-8649.1992.tb02685.x

Knutsen, H., Moland Olsen, E., Ciannelli, L., Espeland, S. H., Knutsen, J. A., Simonsen, J. H., Skreslet, S. and Stenseth, N. C. (2007). Egg distribution, bottom topography and small-scale cod population structure in a coastal marine system. Mar. Ecol. Prog. Ser. 333, 249–255. 10.3354/meps

Knutsen, H., Jorde, P. E., Hutchings, J. A., Hemmer-Hansen, J., Grønkjær, P., Jørgensen, K.-E. M., André, C., Sodeland, M., Albretsen, J. and Olsen, E. M. (2018). Stable coexistence of genetically divergent Atlantic cod ecotypes at multiple spatial scales. Evol. Appl. 11, 1527– 1539. 10.1111/eva.12640

Lüdecke, D. (2018). Ggeffects: Tidy data frames of marginal effects from regression models. J. Open Source Softw. 3, 772. 10.21105/joss.00772

Mangor-Jensen, A. (1987). Water balance in developing eggs of the cod *Gadus morhua* L. Fish Physiol. Biochem. 3, 17–24. 10.1007/BF02183990

Matschiner, M., Barth, J. M. I., Tørresen, O. K., Star, B., Baalsrud, H. T., Brieuc, M. S. O., Pampoulie, C., Bradbury, I., Jakobsen, K. S. and Jentoft, S. (2022). Supergene origin and maintenance in Atlantic cod. *Nat*. Ecol. Evol. 6, 469–481. 10.1038/s41559-022-01661-x

Mérot, C., Oomen, R. A., Tigano, A. and Wellenreuther, M. (2020). A roadmap for understanding the evolutionary significance of structural genomic variation. Trends Ecol. Evol. 35, 561–572. 10.1016/j.tree.2020.03.002

Mousseau, T. A. and Fox, C. W. (1998). The adaptive significance of maternal effects. Trends Ecol. Evol. 13, 403–407. 10.1016/S0169-5347(98)01472-4

Nielsen, E. E., Hemmer-Hansen, J., Larsen, P. F. and Bekkevold, D. (2009). Population genomics of marine fishes: identifying adaptive variation in space and time. Mol. Ecol. 18, 3128–3150. 10.1111/j.1365-294X.2009.04272.x

Nissling, A. and Vallin, L. (1996). The ability of Baltic cod eggs to maintain neutral buoyancy and the opportunity for survival in fluctuating conditions in the Baltic Sea. J. Fish Biol. 48, 217–227. 10.1111/j.1095-8649.1996.tb01114.x

Nissling, A. and Westin, L. (1997). Salinity requirements for successful spawning of Baltic and Belt Sea cod and the potential for cod stock interactions in the Baltic Sea. Mar. Ecol. Prog. Ser. 152, 261–271. 10.3354/meps152261

Nissling, A., Kryvi, H. and Vallin, L. (1994). Variation in egg buoyancy of Baltic cod *Gadus morhua* and its implications for egg survival in prevailing conditions in the Baltic Sea. Mar. Ecol. Prog. Ser. 110, 67–74.

Oomen, R. A. (2019). THE GENOMIC BASIS AND SPATIAL SCALE OF VARIATION IN THERMAL RESPONSES OF ATLANTIC COD (*GADUS MORHUA*).

Oomen, R. A., Juliussen, E., Olsen, E. M., Knutsen, H., Jentoft, S., and Hutchings, J. A. (2021). Cryptic microgeographic variation in responses of larval Atlantic cod to warmer temperatures. bioRxiv. 10.1101/2021.02.03.429645

Oomen, R. A., Knutsen, H., Olsen, E. M., Jentoft, S., Stenseth, N. C. and Hutchings, J. A. (2022). Warming accelerates the onset of the molecular stress response and increases mortality of larval Atlantic cod. Integr. Comp. Biol. 62, 1784–1801. 10.1093/icb/icac145

Pampoulie, C., Berg, P. R. and Jentoft, S. (2023). Hidden but revealed: After years of genetic studies behavioural monitoring combined with genomics uncover new insight into the population dynamics of Atlantic cod in Icelandic waters. Evol. Appl. 16, 223–233. 10.1111/eva.13471

R Core Team. (2023). R: A Language and Environment for Statistical Computing. R Foundation for Statistical Computing, Vienna, Austria. <https://www.R-project.org/>.

Roney, N. E., Oomen, R. A., Knutsen, H., Olsen, E. M., and Hutchings, J. A. (2017). Temporal variability in offspring quality and individual reproductive output in a broadcast-spawning marine fish. ICES J. Mar. Sci. 75, 1353–1361. 10.1093/icesjms/fsx232

Saborido-Rey, F. (2003). Buoyancy of Atlantic cod larvae in relation to developmental stage and maternal influences. J. Plankton Res. 25, 291–307. 10.1093/plankt/25.3.291

Schmidt, N., Garate-Olaizola, M. and Laurila, A. (2024). Acclimatizing laboratory-reared hatchling cod (*Gadus morhua*) to salinity conditions in the Baltic Sea. Aquaculture 579, 740255. 10.1016/j.aquaculture.2023.740255

Shapero, M. (2013). SNP Genotyping using Affymetrix’ Axiom® Genotyping Solution.

Sinclair-Waters, M., Bradbury, I. R., Morris, C. J., Lien, S., Kent, M. P. and Bentzen, P. (2018). Ancient chromosomal rearrangement associated with local adaptation of a postglacially colonized population of Atlantic Cod in the northwest Atlantic. Mol. Ecol. 27, 339–351. 10.1111/mec.14442

Sodeland, M., Jorde, P. E., Lien, S., Jentoft, S., Berg, P. R., Grove, H., Kent, M. P., Arnyasi, M., Olsen, E. M. and Knutsen, H. (2016). “Islands of divergence” in the Atlantic cod genome represent polymorphic chromosomal rearrangements. Genome Biol. Evol. 8, 1012–1022. 10.1093/gbe/evw057

Sodeland, M., Jentoft, S., Jorde, P. E., Mattingsdal, M., Albretsen, J., Kleiven, A. R., Synnes, A.-E. W., Espeland, S. H., Olsen, E. M., Andrè, C., et al. (2022). Stabilizing selection on Atlantic cod supergenes through a millennium of extensive exploitation. Proc. Natl. Acad. Sci. U. S. A. 119, e2114904119. 10.1073/pnas.2114904119

Stenevik, E. K., Sundby, S. and Agnalt, A. L. (2008). Buoyancy and vertical distribution of Norwegian coastal cod (*Gadus morhua*) eggs from different areas along the coast. ICES J. Mar. Sci. 65, 1198–1202. 10.1093/icesjms/fsn101

Sundby, S., and Kristiansen, T. (2015). The principles of buoyancy in marine fish eggs and their vertical distributions across the world oceans. PloS one 10(10), e0138821. 10.1371/journal.pone.0138821

Thomas, P. D., Ebert, D., Muruganujan, A., Mushayahama, T., Albou, L. P., and Mi, H. (2022). PANTHER: Making genome-scale phylogenetics accessible to all. Protein Sci. 31(1), 8–22. 10.1002/pro.4218

Wellenreuther, M., Oomen, R. A., Young Han, K., Krohman, R., and T. B. H. Reusch. (2025). Beyond supergenes: the diverse roles of inversions in trait evolution. Trends Ecol. Evol. ISSN 0169–5347. 10.1016/j.tree.2025.08.004

Whitlock M. C. and Lotterhos K. (2014). OutFLANK: Fst outliers with trimming. R package version 0.2. https://github.com/whitlock/OutFLANK/ (last accessed 12 January 2022).

Wickham, H. (2016). ggplot2: Elegant Graphics for Data Analysis. Springer-Verlag New York.

Wolf, J. B. and Wade, M. J. (2009). What are maternal effects (and what are they not)? Philos. Trans. R. Soc. Lond. B Biol. Sci. 364, 1107–1115. 10.1098/rstb.2008.0238

