## Supplementary Notes for "Atlantic cod (Gadus morhua) ecotypes, not inversion frequencies, underlie divergence in egg buoyancy distribution"

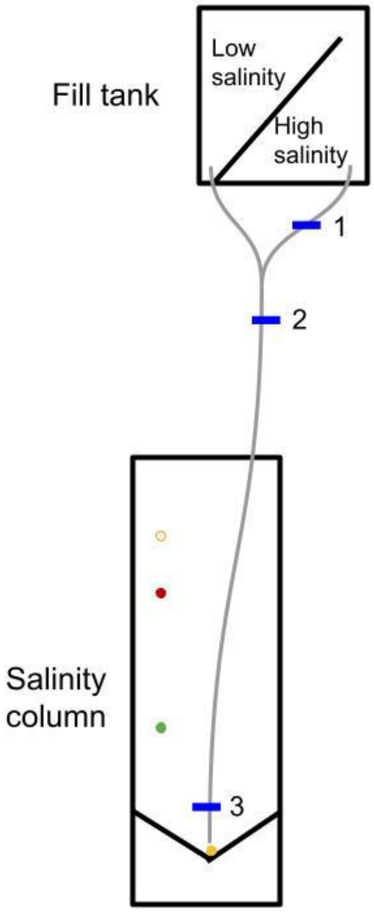


**Figure S1.** A schematic of the salinity gradient system. In the drawn diagram, the fill tank and salinity column are black, clamps 1-3 are blue and labelled, and the tubing is gray. The four circles in the schematic are the four beads in their respective colours and located in the salinity column around the location where they would sit.

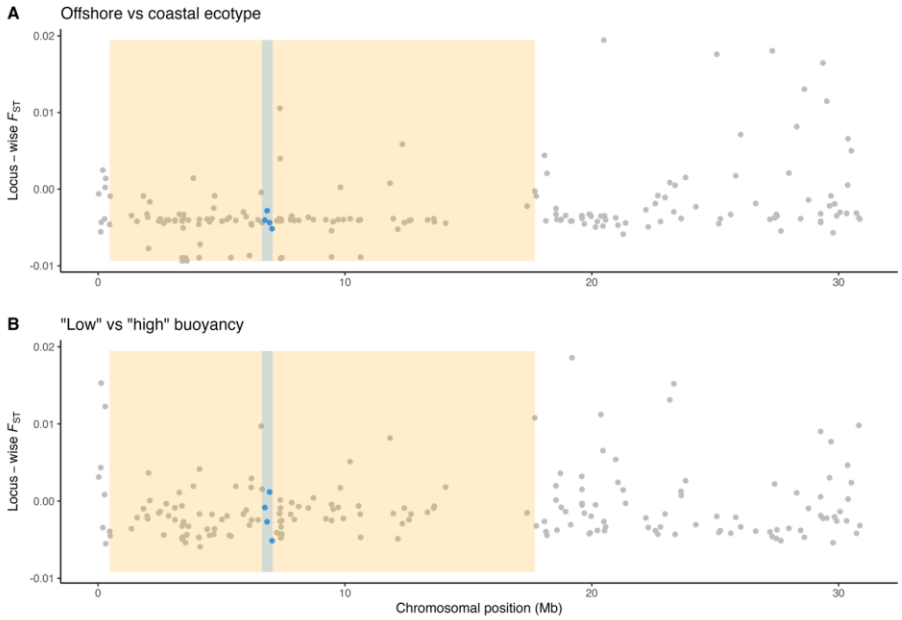


**Figure S2.** Pairwise $F_{ST}$ along chromosome 12 between A) North Sea and Fjord ecotype eggs, and B) "low" and "high" buoyancy. The inverted region is highlighted in orange, the region of the double crossover is highlighted in blue, while loci therein are indicated with blue points.

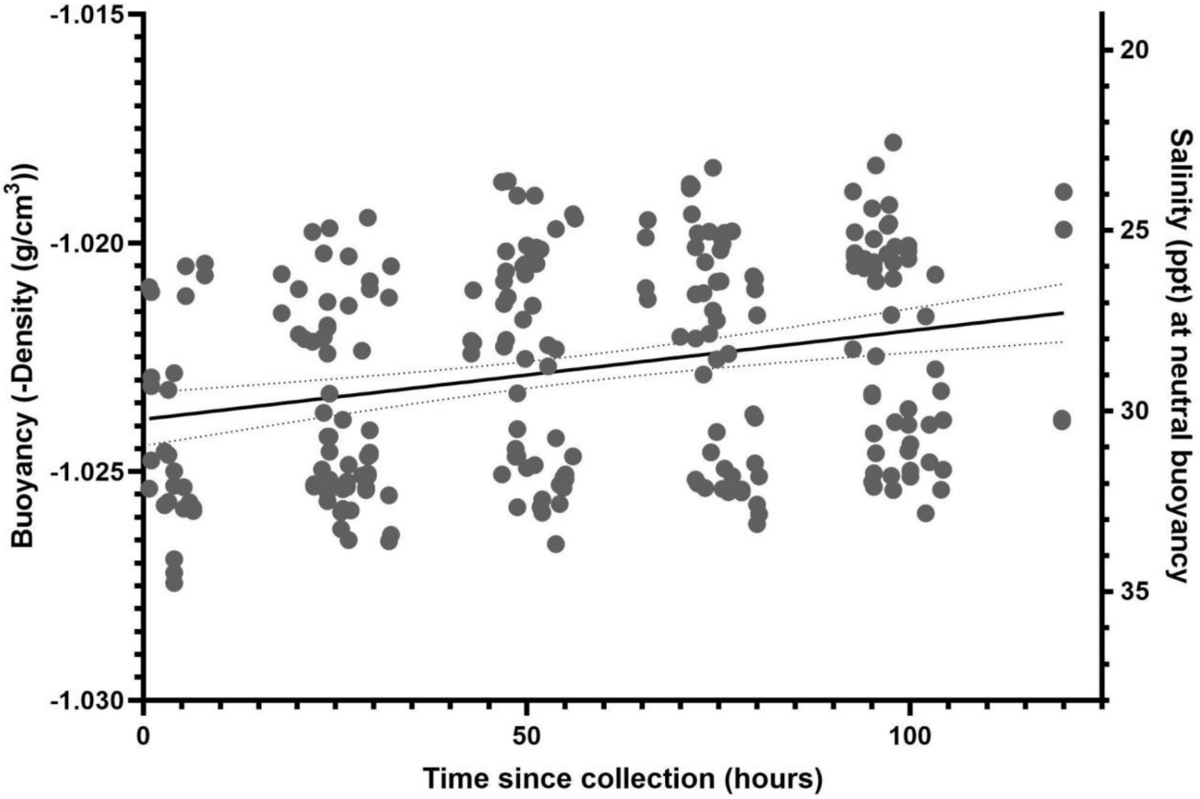


**Figure S3.** The buoyancy (-Density ($g/cm^3$)) of all eggs increases with the time since collection (hours). The neutral buoyancy at a given salinity (ppt) at 7°C is provided to help link salinity with buoyancy. Plotted points are the raw data points. Dotted lines show the 2.5% to 97.5% confidence intervals for the line of best fit. N=250.

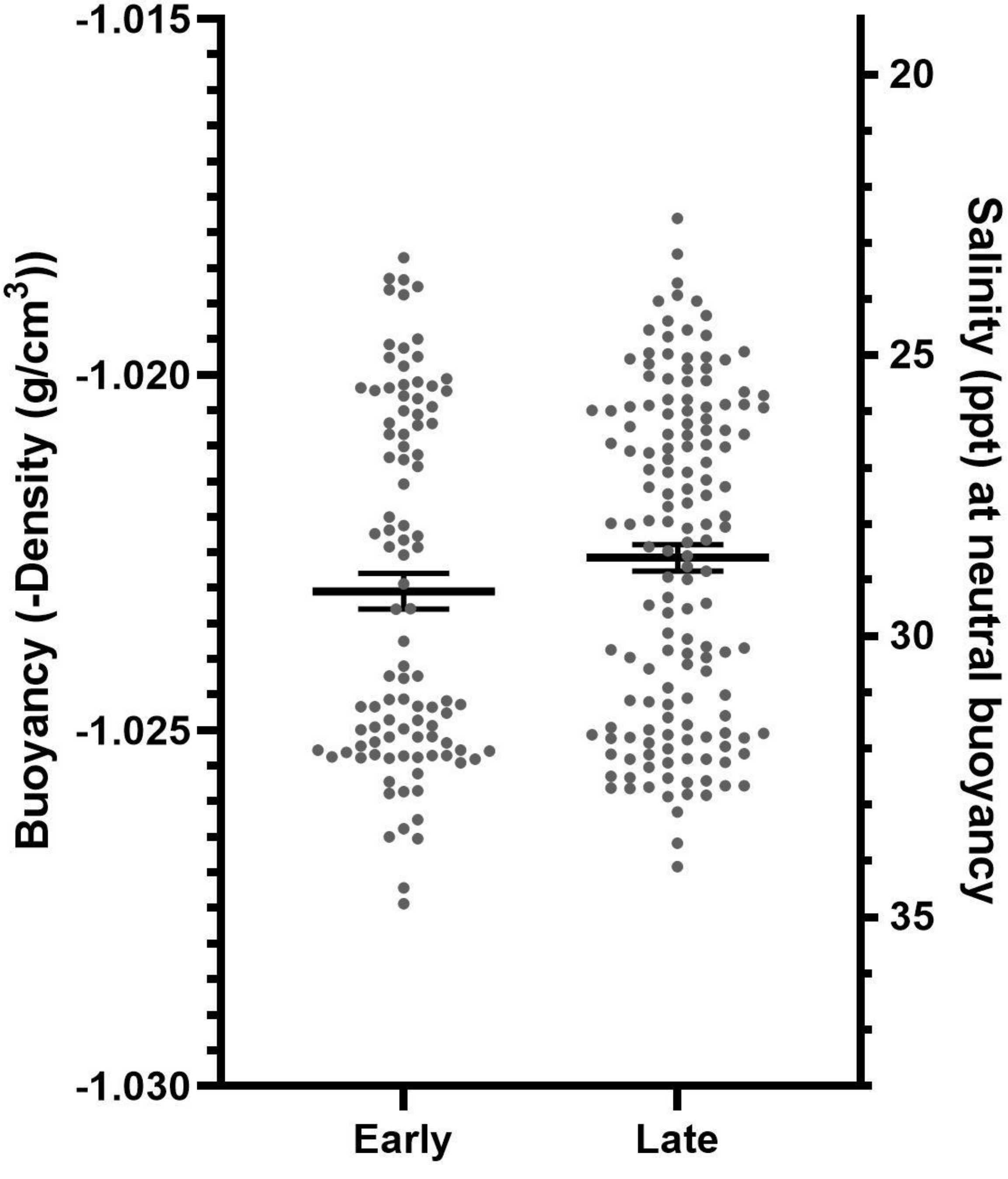


**Figure S4.** There is little difference in buoyancy (-Density ($g/cm^3$)) between early and late egg stages. The neutral buoyancy at a given salinity (ppt) at 7°C is provided to help link salinity with buoyancy. Plotted points are the raw data points. Error bars represent the standard error around the mean. n = 99 and 151 for early and late individuals, respectively.

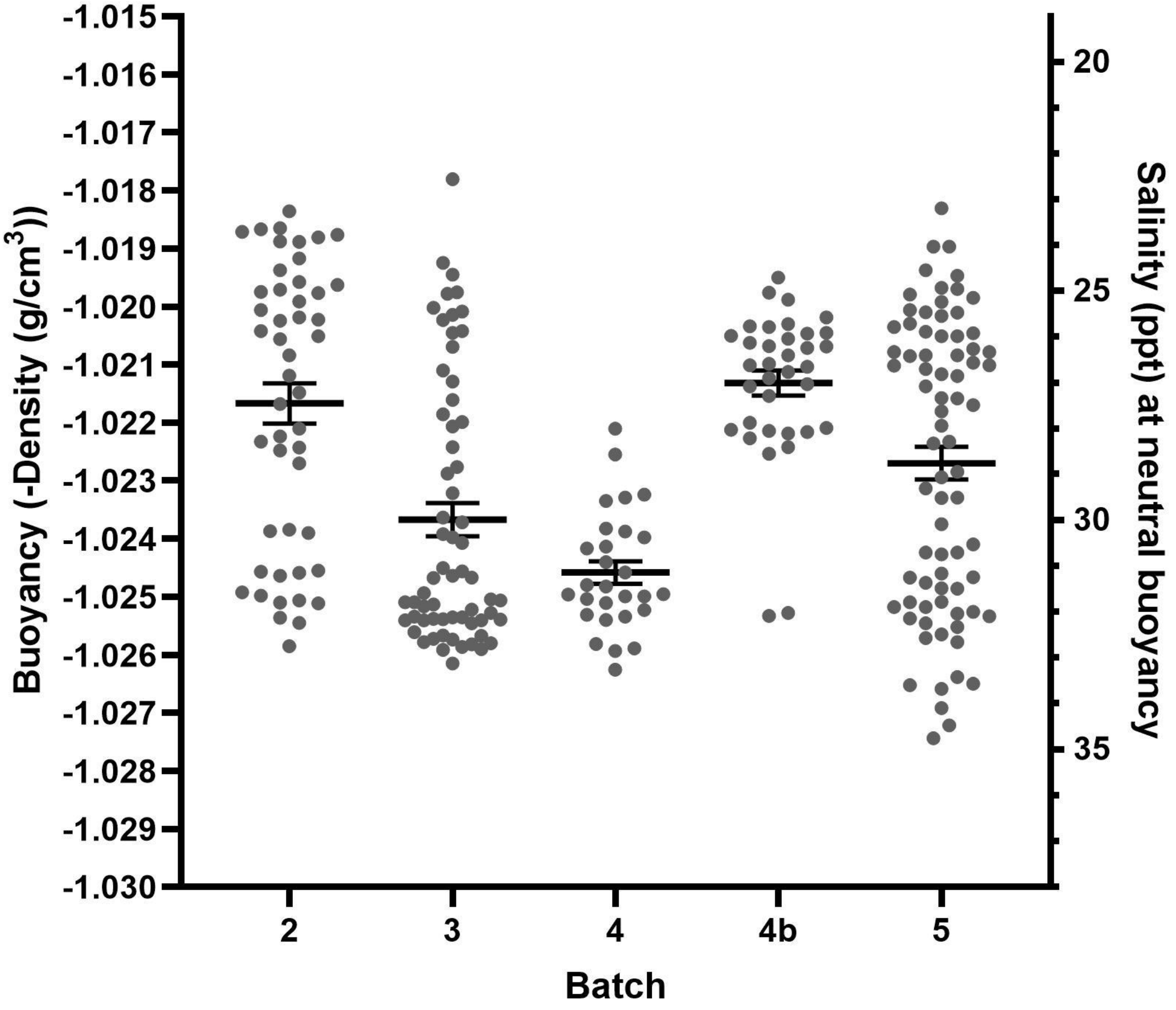


**Figure S5.** Egg buoyancy (-Density (g/cm$^3$)) differs among batches. The neutral buoyancy at a given salinity (ppt) at 7°C is provided to help link salinity with buoyancy. Plotted points are the raw data points. Error bars represent the standard error around the mean. n=48, 63, 28, 35, and 76 for batches 2, 3, 4, 4b, and 5, respectively.

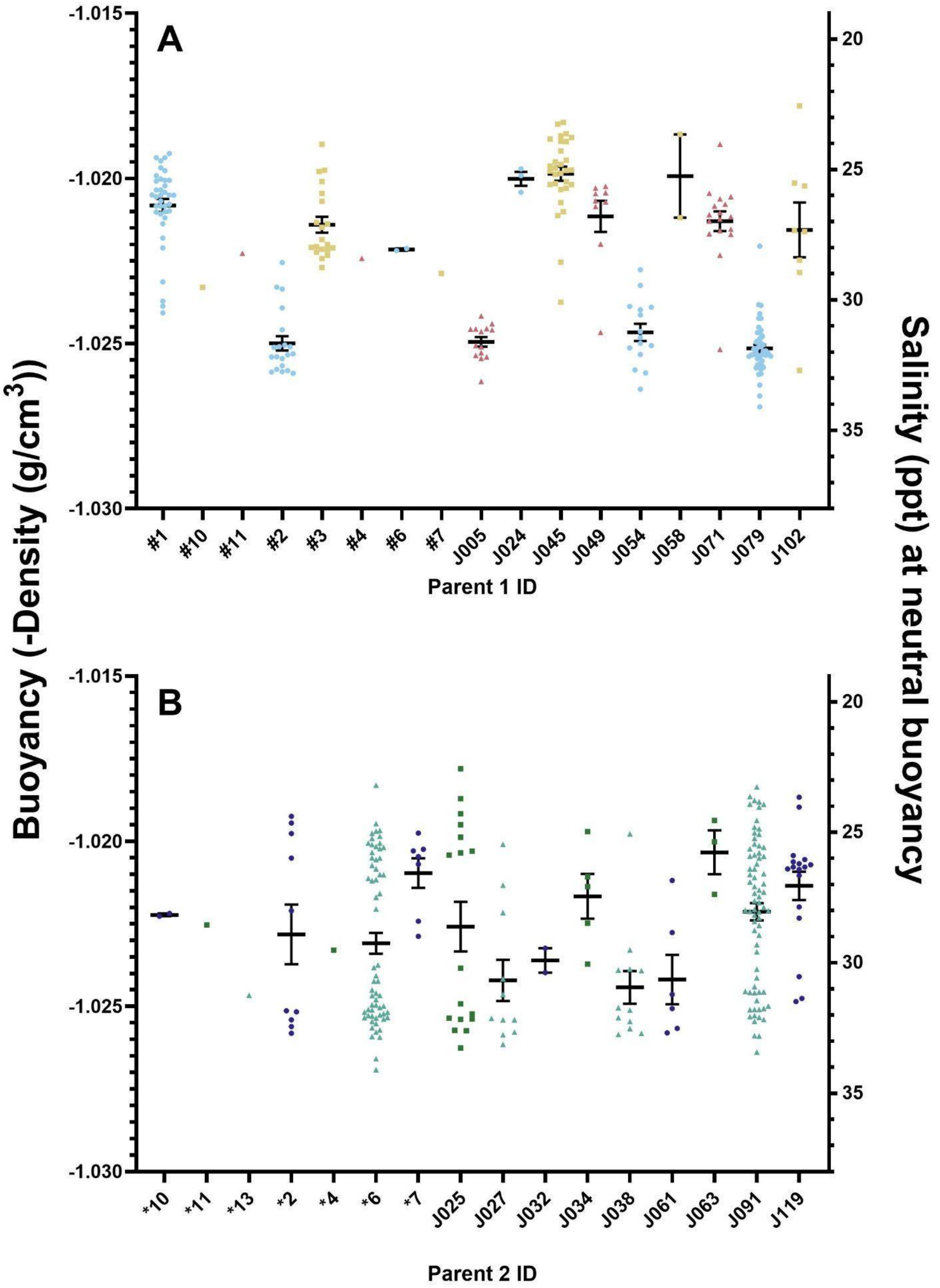


**Figure S6.** Egg buoyancy differed between parents 1 (A) and 2 (B). Plotted points are the raw data points. Error bars represent the standard error around the mean. N=230.

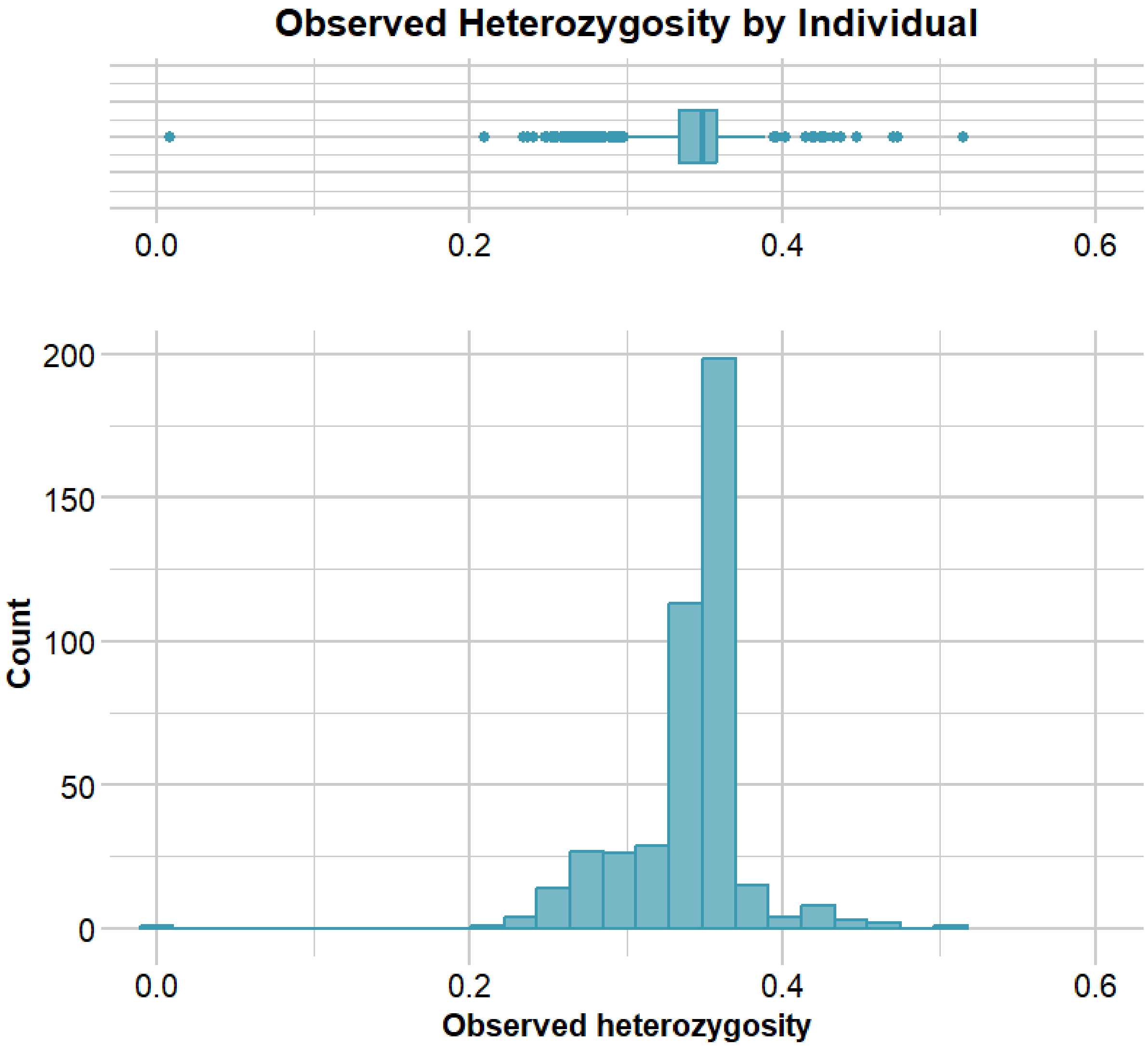


**Figure S7.** A histogram and box plot showing the observed heterozygosity for all 446 samples sent off for DNA analysis.

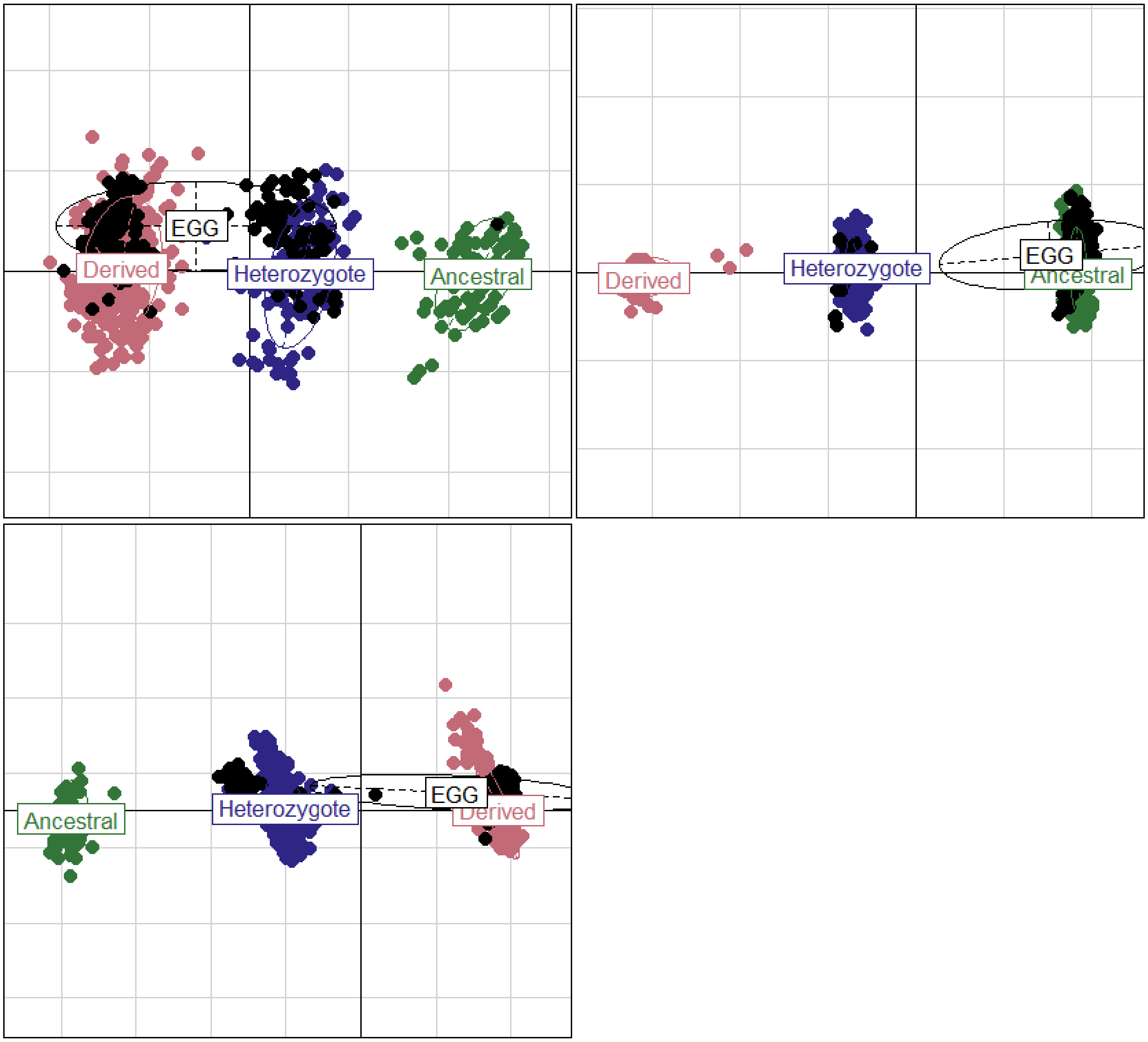


**Figure S8.** PCA showing the genotype classifications for inv2 (A), inv7 (B), and inv12 (C) for all the individuals from the baseline dataset and where the individuals in the egg dataset are located on the PCAs.

**Table S1.** The amount of variance explained by each random effect in the model.

| Random effect | Variance explained |
| --- | --- |
| Stage | 5.140e-15 |
| Parent 1 ID | 2.875e-06 |
| Parent 2 ID | 6.060e-07 |
| Time since collection | 6.375e-07 |
| Batch | 7.221e-07 |
