## Supplementary Methods for "Atlantic cod (Gadus morhua) ecotypes, not inversion frequencies, underlie divergence in egg buoyancy distribution"

### 1 Collection of batch 3 and 4

In batch 3, water from the laboratory was used in the collection bucket (instead of surface pond water) due to poor surface water conditions. For batch 4, two collections, instead of one, took place according to the previously stated methods, 75 minutes apart. However, few individuals were collected in these two collections. There were thousands of eggs visible at the surface of the northeast corner of the pond, likely due to a strong northeast wind. These eggs were collected (3rd collection for batch 4) with a 1 L plastic container 5 hours and 45 minutes after the first collection. These individuals were termed batch 4b and were kept in holding container I (Figure 2), separated from batch 4. Batch 4b was included in the analyses to ensure that the variation of individuals in the pond at that time was captured. Only individuals with lower buoyancies likely avoided getting blown to the corner of the pond because they would sit below the surface at neutral buoyancy. In the wild, wind would create waves that would mix eggs into the ocean rather than blow them into one area, and a sheltered area would not likely be impacted by strong winds.

### 2 Assigning inversion genotypes, ecotypes, and parental relationships

All analyses used to classify SNP data were completed in R v4.4.2 using R Studio (R Core Team, 2023). Figures were created using the ggplot2 package in R (v3.5.2; Wickham, 2016)

A dataset including 466 cod samples, created using the same SNP panel, including east Atlantic North Sea, east Atlantic Fjord, and eastern Baltic cod, was used as a baseline dataset to help assign the inversion genotypes to the eggs (Henriksson et al., in prep.; *cf.* Andersson et al., 2023). First, we filtered the 3776 loci contained within the baseline data to only include loci found in our dataset (3462 total loci). Our dataset was double-checked to make sure that all individuals contained SNP data, and we combined the two datasets manually in a text file (hereafter "combined data"). After, the loci in the locus metadata were lined up with the loci in the combined data and loci not included in the combined data were removed from the cod locus metadata. The call rate per locus and individual was tested in the egg data and was determined to be above 0.80 (90.134% min call rate for loci, 95.07% min call rate for individuals), suggesting no loci or individuals needed to be removed due to poor quality genotyping. Heterozygosity was used to test for contamination in the dataset. The gl.report.heterozygosity function in the dartR package (v2.9.7; Gruber et al., 2018) was used to calculate heterozygosity per individual. A mean and median heterozygosity of 0.34 and 0.35 was found, respectively (Figure S7). There were no extremities seen in the heterozygosity, indicating that contamination is unlikely.

Principal Component Analysis (PCA) based on sequenced SNPs was used to identify the inversion genotypes on chromosomes 2, 7, and 12. We created a PCA for inv2, inv7, and inv12 using the baseline dataset with loci found in inv2 (18 loci), inv7 (74 loci), and inv12 (101 loci), using the dudi.pca function in the ade4 package (v1.7.22; Dray and Dufour, 2007). Combined data were plotted onto these PCAs, one individual at a time, to account for kinship. PCA axes were oriented in the same direction for each plotting of an individual. To orient the axes, we coded 4 individuals with known locations to be in the same locations relative to each other in each iteration. We then assigned inversion genotypes to egg individuals based on the location of the individual's data point relative to the baseline dataset for each genotype (Figure S8).

The individuals in the baseline dataset also acted as a reference for ecotype assignment. The percent likelihood that each individual in the baseline dataset and the combined dataset belonged to each ecotype was determined by comparing individuals to the baseline dataset using the assign.X function in the assignPOP package (v1.3.0; Chen, 2024). The maximum percent likelihood that any egg belonged to the eastern Baltic ecotype was 0.01925, and the mean±SE was 0.005903±0.000167, meaning all eggs likely belonged to the Fjord or North Sea ecotypes. We did not classify individuals as ecotype factors (ex., North Sea, Fjord, and hybrids) because there were no clear cutoffs in the percent likelihood of belonging to a given ecotype.

We determined parent-offspring relationships using the programme COLONY (v2.0.7.1; Wang, 2004). A dataset with 120 cod from the parental generation (F1 generation), genotyped using the same SNP panel (Henriksson et al., in prep), was used as potential parents for the eggs collected in this experiment. The full-likelihood method was used with high precision and random seed numbers. Genotyping error was set to 0.0001 per locus. We repeated the analysis four times using a medium-length run each time. The runs were combined manually, and the results of the runs were matched.

**3 Model results with random effects**

We tested the effect of each random effect, egg stage, parent 1 and 2 ID, time since collection, and batch, on the model individually (ex., Egg buoyancy ~ inv2 + inv7 + inv12 + probability of belonging to the North Sea ecotype + [1| batch]). We found that each random effect explained minimal variance in the model (Table S1). This was likely due to there being correlations between our main effects and random effects.
