## Supplementary figure and Tables for "Atlantic cod (Gadus morhua) ecotypes, not inversion frequencies, underlie divergence in egg buoyancy distribution"

### Supplementary Notes

#### Genomic differentiation

Outlier analysis detected five outlier loci between North Sea and Fjord ecotype eggs, located within CLEC16A, PTPRE, GRID1, ADSS2, and ASTN2 genes. CLEC16A (C-Type Lectin Domain Containing 16A) is suggested to play a role in neurogeneration/-degeneration, immune response, and mitochondrial health in zebrafish (*Danio rerio*; Smits et al., 2023). PTPRE (Protein Tyrosine Phosphatase Receptor Type E) is shown to impact cell movement (van Eekelen et al., 2012), and gastrulation in zebrafish (van Eekelen et al., 2010), and growth in both zebrafish (van Eekelen et al., 2012) and tambaqui (*Colossoma macropomum*; Ariede et al., 2023). GRID1 (Glutamate Ionotropic Receptor Delta Type Subunit 1), similarly, is linked with growth in tongue sole (*Cynoglossus semilaevis*; Wang et al., 2025), but also with environmental adaptation in several fish species (Kim et al., 2016; Zhou et al., 2023; Zoghly et al., 2023). ADSS2 (Adenylosuccinate Synthase 2) plays a role in nucleotide metabolism in northern snakehead (*Channa argus*; Sun et al., 2025), but also reproductive maturity in both channel catfish (*Ictalurus punctatus*; Li et al., 2017) and common carp (*Cyprinus carpio*; Su et al., 2014). Finally, ASTN2 (Astrotactin 2) likely plays a role in muscle filament assembly in tongue sole (Lü et al., 2025).

The outlier analysis between "low" and "high" buoyancy eggs found three loci, two of which were located within genes (MACROD2 and GBF1), while one locus was located between two genes – 126 bases downstream of RRM1 and 524 bases upstream of SIPA1l3. MACROD2 (Mono-ADP Ribosylhydrolase 2) has GO terms associated with protein modification, nucleoside metabolism, and DNA damage response, but has not yet, to our knowledge, been studied in fish. GBF1 (Golgi Brefeldin A Resistant Guanine Nucleotide Exchange Factor 1) is been linked with body growth rate in Nile tilapia (*Oreochromis niloticus*; Zhang et al., 2022), and fin shape and development in both zebrafish (Chen et al., 2017) and *Lamprologini* cichlids (Ahi et al., 2022). RRM1 (Ribonucleotide Reductase Catalytic Subunit M1), located upstream of the outlier locus on chromosome 16, is linked to DNA damage response in zebrafish (Danilova et al., 2014) and response to viral infection in both Jinhu grouper (*Epinephelus fuscoguttatus*♀ × *Epinephelus tukula*♂; Duan et al., 2025) and Asian bass (*Lates calcarifer*; Yang et al., 2021). Lastly, SIPA1l3 (Signal Induced Proliferation Associated 1 Like 3), located 524 bases downstream of the outlier locus on chromosome 16, is shown to play an important role in electrical synapse function in zebrafish (Tetenborg et al., 2024).

#### References for Supplementary Notes

**Ahi, E. P., Richter, F., and Sefc, K. M.** (2022). Gene expression patterns associated with fin shape differ between two lamprologine cichlids. *bioRxiv*, 2022-06.

**Chen, J., Wu, X., Yao, L., Yan, L., Zhang, L., Qiu, J., ... and Meng, A.** (2017). Impairment of cargo transportation caused by gbf1 mutation disrupts vascular integrity and causes hemorrhage in zebrafish embryos. *Journal of Biological Chemistry*, **292**(6), 2315-2327.

**Danilova, N., Bibikova, E., Covey, T. M., Nathanson, D., Dimitrova, E., Konto, Y., ... and Lin, S.** (2014). The role of the DNA damage response in zebrafish and cellular models of Diamond Blackfan anemia. *Disease models & mechanisms*, **7**(7), 895-905.

**Duan, H., Tian, Y., and Li, Z.** (2025). Transcriptome research conducted on the liver and spleen of Jinhu grouper (Epinephelus fuscoguttatus♀× Epinephelus tukula♂) reveals the mechanism in response to Vibrio anguillarum infection. *Comparative Biochemistry and Physiology Part D: Genomics and Proteomics*, **55**, 101482.

**Kim, Y. J., Lee, N., Woo, S., Ryu, J. C., and Yum, S.** (2016). Transcriptomic change as evidence for cadmium-induced endocrine disruption in marine fish model of medaka, Oryzias javanicus. *Molecular & Cellular Toxicology*, **12**(4), 409-420.

**Li, H., Su, B., Qin, G., Ye, Z., Alsaqufi, A., Perera, D. A., ... and Dunham, R. A.** (2017). Salt sensitive Tet-off-like systems to knockdown primordial germ cell genes for repressible transgenic sterilization in channel catfish, Ictalurus punctatus. *Marine Drugs*, **15**(6), 155.

**Lü, Z., Wang, Y., Yu, J., Yang, Y., Xu, A., Gong, L., ... and Liu, L.** (2025). Comparison of muscle structure and transcriptome analysis of eyed-side muscle and blind-side muscle in Cynoglossussemilaevis (Osteichthyes, Cynoglossidae). *ZooKeys*, **1230**, 213.

Raquel B. Ariede, Milena V. Freitas, Rubens R. Oliveira Neto et al. Genome wide association study for growth and carcass traits in the Amazon fish Colossoma macropomum, 20 December 2023, PREPRINT (Version 1) available at Research Square. https://doi.org/10.21203/rs.3.rs-3750262/v1

**Smits, D. J., Dekker, J., Schot, R., Tabarki, B., Alhashem, A., Demmers, J. A., ... and Mancini, G. M.** (2023). CLEC16A interacts with retromer and TRIM27, and its loss impairs endosomal trafficking and neurodevelopment. Human Genetics, **142**(3), 379-397.

**Su, B., Peatman, E., Shang, M., Thresher, R., Grewe, P., Patil, J., ... and Dunham, R. A.** (2014). Expression and knockdown of primordial germ cell genes, vasa, nanos and dead end in common carp (Cyprinus carpio) embryos for transgenic sterilization and reduced sexual maturity. *Aquaculture*, **420**, S72-S84.

**Sun, D., Tao, Z., Wen, H., Qi, X., Li, C., Wang, L., ... and Li, Y.** (2025). Comparative analysis of muscle quality, transcriptomic, and metabolomic profiles in yellow-mutant and wild-type northern snakehead (Channa argus): Implications for food quality and aquaculture breeding potential. *Food Bioscience*, **68**, 106496.

**Tetenborg, S., Shihabeddin, E., Kumar, E. O. A. M., Sigulinsky, C. L., Dedek, K., Lin, Y. P., ... and O'Brien, J.** (2024). Uncovering the electrical synapse proteome in retinal neurons via in vivo proximity labeling. *bioRxiv*, 2024-11.

**van Eekelen, M., Runtuwene, V., Overvoorde, J., and den Hertog, J.** (2010). RPTPα and PTPε signaling via Fyn/Yes and RhoA is essential for zebrafish convergence and extension cell movements during gastrulation. Developmental biology, **340**(2), 626-639.

**van Eekelen, M., Runtuwene, V., Masselink, W., and den Hertog, J.** (2012). Pair-wise regulation of convergence and extension cell movements by four phosphatases via RhoA. PLoS One, **7**(4), e35913.

**Wang, Y., Chen, Y., Liu, Y., and Chen, S.** (2025). Molecular Mechanism of the Grid Gene Family Regulating Growth Size Heteromorphism in Cynoglossus semilaevis. Animals, **15**(8), 1130.

**Yang, Z., Wong, S. M., and Yue, G. H.** (2021). Effects of rrm1 on NNV resistance revealed by RNA-seq and gene editing. *Marine Biotechnology*, **23**(6), 854-869.

**Zhang, H., Rambhau, J. N., Chae, J. Y., Yoo, B. H., and Lee, H. H.** (2022). Relationship between single nucleotide polymorphism of IGF-1 gene in Nile tilapia (Oreochromis niloticus) and weight gain. *Authorea Preprints*.

**Zhou, Z., Yang, J., Lv, H., Zhou, T., Zhao, J., Bai, H., ... and Xu, P.** (2023). The adaptive evolution of Leuciscus waleckii in Lake Dali Nur and convergent evolution of Cypriniformes fishes inhabiting extremely alkaline environments. *Genome Biology and Evolution*, **15**(5), evad082.

**Zoghly, H., Rashed, M., and Magdy, M.** (2023). Partial Gap-Filling of the Nile Tilapia Oreochromis niloticus Draft Genome. *Arab Universities Journal of Agricultural Sciences*, **31**(1), 119-130.
